# Characterization and establishment of a novel EBV strain simultaneously associated with nasopharyngeal carcinoma and B-cell lymphoma

**DOI:** 10.1101/2020.03.26.009894

**Authors:** Fenggang Yu, Nicholas L. Syn, Yanan Lu, Qing Yun Chong, Junyun Lai, Wei Jian Tan, Boon Cher Goh, Paul A. MacAry, Lingzhi Wang, Kwok Seng Loh

**Author notes:** Equal contributions (Fenggang Yu, Nicholas L. Syn). Cancer Immunology Program, Peter MacCallum Cancer Centre, Melbourne, VIC 3000, Australia. Sir Peter MacCallum Department of Oncology, The University of Melbourne, Parkville, VIC 3000, Australia. **Correspondence:** Fenggang Yu, PhD; Department of Otolaryngology, National University Health System, Singapore; 1E Kent Ridge Road, Level 7, NUHS Tower Block, Singapore 119228. **Disclosures:** The authors declare no competing interests.

## Abstract

Epstein-Barr virus (EBV) – the prototypical human tumor virus – is responsible for 1-2% of the global cancer burden, but divergent strains seem to exist in different geographical regions with distinct predilections for causing lymphoid or epithelial malignancies. Here we report the establishment and characterization of Yu103, an Asia Pacific EBV strain with a highly remarkable provenance of being derived from nasopharyngeal carcinoma biopsy but subsequently propagated in human B-lymphoma cells and xenograft models. Unlike previously characterized EBV strains which are either predominantly B-lymphotropic or epitheliotropic, Yu103 evinces an uncanny capacity to infect and transform both B-lymphocytes and nasopharyngeal epithelial cells. Genomic and phylogenetic analyses indicated that Yu103 EBV lies midway along the spectrum of EBV strains known to drive lymphomagenesis or carcinogenesis, and harbors molecular features which likely account for its unusual properties. To our knowledge, Yu103 EBV is currently the only EBV isolate shown to drive human nasopharyngeal carcinoma and B-lymphoma, and should therefore provide a powerful novel platform for research on EBV-driven hematological and epithelial malignancies.

## INTRODUCTION

Epstein-Barr virus (EBV), the prototypical human tumor virus [1], infects 95% of the human adult population and is etiologically implicated in a spectrum of lymphoid and epithelial malignancies, which exhibit very striking patterns of worldwide distribution. Whereas Hodgkin’s lymphomas feature a much higher prevalence in Europe and North America, Burkitt’s lymphoma is endemic in equatorial Africa, and non-keratinizing nasopharyngeal carcinoma (NPC) in East and Southeast Asia is almost universally associated with EBV infection [2,3]. This epidemiological enigma, as well as the observation that only a small minority of individuals infected with the virus actually develop neoplastic disease during their lifetimes, has led to the hypothesis that different strains of EBV prevail worldwide with varying propensities for infecting and subsequently transforming B cells and epithelial cells. Indeed, although most EBV strains characterized in the context of malignancy are predominantly B-lymphotropic [4–7], the few strains that have been studied in NPC have been found to harbor unique genetic polymorphisms in certain viral sequences (e.g. *LMP1* and *BZLF1* promoter) which endow them with greatly enhanced epitheliotropism [5,8,9]. Distinct molecular processes appear to be involved in orchestrating the virus’ attachment, entry into, and replication in B lymphocytes and epithelial cells [4,6,10–12].

Hitherto, preclinical investigations into the role and therapeutic implications of EBV in NPC, a unique epithelial malignancy with a median survival of 12-20 months in the recurrent and metastatic settings [2,3,13], have almost exclusively relied on the C666-1 cell line – which until recently, was the only extensively-used EBV-positive NPC cell line in research [9]. The scarcity of *in vitro* NPC models which stably maintain the viral episome has invariably hampered translational and therapeutic advances, including novel immunotherapies (eg. adoptive T-cell therapies and therapeutic cancer vaccines directed against EBV epitopes) and prophylactic vaccines whose development are contingent on the ability of preclinical models to recapitulate the repertoire and variation in viral epitopes across different EBV strains [2,14–16].

Here we report the establishment and characterization of Yu103, an Asian Pacific EBV strain that has a highly unusual provenance of being derived from a nasopharyngeal carcinoma biopsy but subsequently propagated in a B cell lymphoma cell line. Unlike most previously characterized EBV strains that are either predominantly B-lymphotropic or epitheliotropic, Yu103 demonstrates an innately programmed capacity to not only infect, but also transform, both B lymphocytes and nasopharyngeal epithelial cells, and should therefore provide a powerful novel platform for research on EBV-associated hematological and epithelial malignancies.

## RESULTS

### Establishment of patient-derived xenograft (PDX) from nasopharyngeal carcinoma biopsy in immunodeficient mice

A biopsy was obtained from a 38-year-old Singaporean Chinese male with stage 4 (T3N1M0) nasopharyngeal carcinoma. Enzymatically-dissociated cells began to proliferate after plating and achieved confluence within a week. The majority of cells exhibited a typical polygonal, cuboidal epithelial appearance, occasionally seen along with spindle-shaped fibroblasts and floating small round cells loosely attached to the well. NOD-scid IL2rynull (NSG) mice were subcutaneously implanted with trypsinized cells, and the xenograft tumor was resected at a diameter of ~15mm at approximately 3 weeks after transplantation (**Figure 1a**).

**Figure 1.**
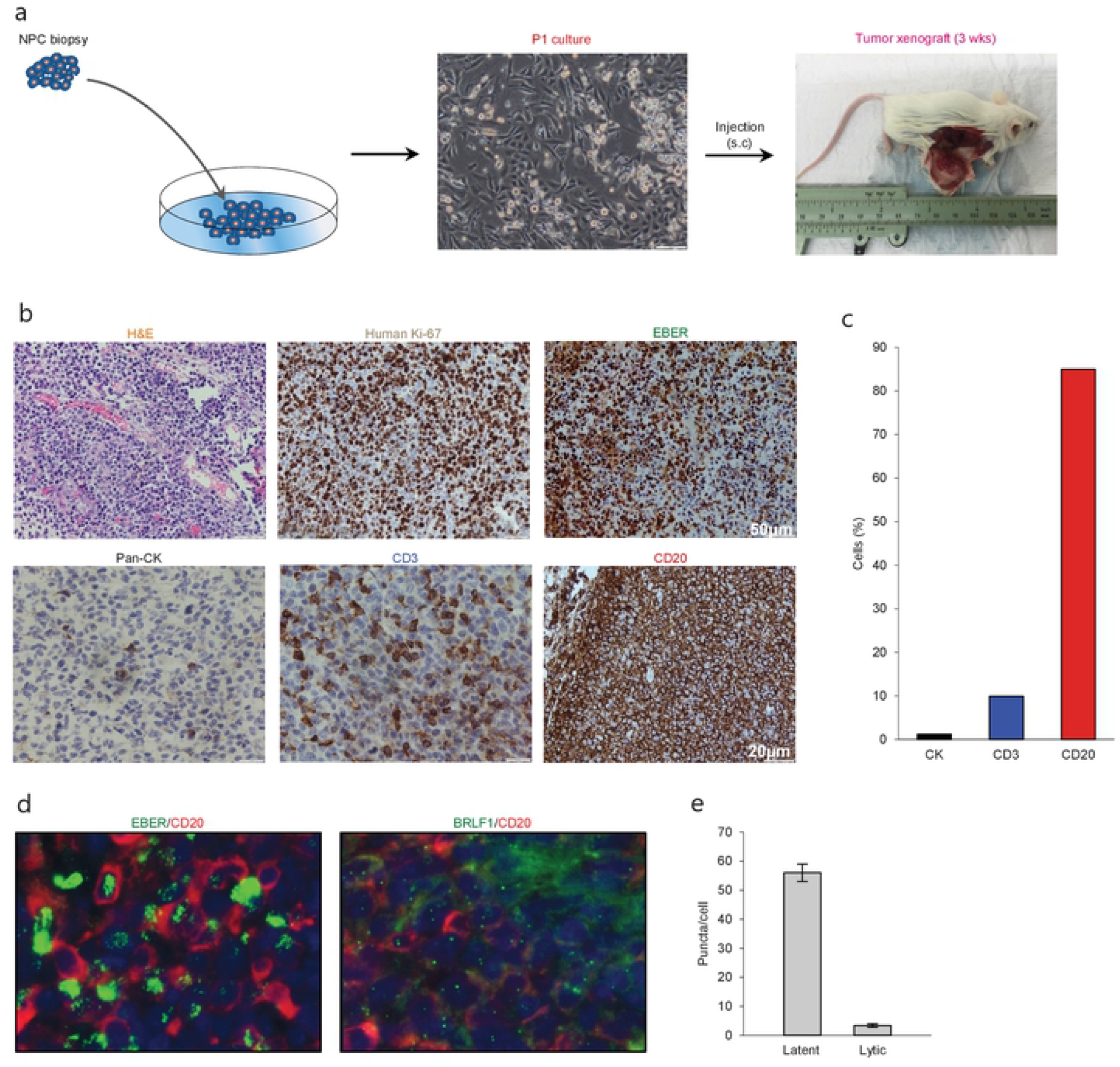
Histological characterization of P1 culture and parental xenograft. (a) A biopsy was obtained from a 38-year-old Singaporean Chinese male with nasopharyngeal carcinoma. The majority of cells in the P1 culture exhibited a typical polygonal, cuboidal epithelial appearance, occasionally seen along with spindle-shaped fibroblasts and floating small round cells loosely attached to the well. NOD-scid IL2rγnull (NSG) mice were subcutaneously implanted with trypsinized cells, and the xenograft tumor was harvested at a diameter of ~15mm at approximately 3 weeks after transplantation. (b) Analysis of histology (H&E), species origin and mitotic rate (human specific Ki67), EBV status (EBER), and tissue composition (lineage markers: pan-CK, CD3, and CD20). (c) Bar graph showing the composition of cell types by lineage markers of the tumor xenograft. (d-e) 4.0μm tissue sections were in in situ hybridized with EBV latent- and lytic-specific RNA probes EBER and BRLF1 respectively, followed by immunohistochemical staining with the B-lymphocyte marker CD20. Positive ISH signals were localized within the cell nuclei as green fluorescent puncta, while CD20 was localized on the cell surface.

### Histopathological characterization of PDX revealed predominance of EBV+ human B cell lymphoma cells

Hematoxylin- and eosin-stained 4-μm sections of the resected xenograft revealed an abundance of small round cells with darkly-stained nucleus. Using a human-specific anti-ki67 antibody, we confirmed that the xenograft was composed of human cells with high mitotic index and not murine cells. Further analysis by cell lineage markers revealed that the majority of cells expressed CD20 (>85%), which is indicative of B cell lineage (**Figure 1A**), with smaller populations of cells that expressed pan-cytokeratin (<5%) or CD3 (<10%) (**Figure 1B**). To establish the presence of EBV and to characterize its replication cycle in the CD20+ subpopulation, we exploited chromogenic EBER-ISH as well as fluorescent EBER-ISH and BRLF-ISH assays, which detect gene products specific to latency and lytic transcriptional programs[17,18]. These assays revealed that although latent infection was predominant, the virus also underwent spontaneous lytic reactivation in B cells at unusually high levels (**Figures 1A and 1C**). Collectively, these findings demonstrated that the xenograft was primarily composed of an aggressive form of EBV-associated human B-cell lymphoma.

The serendipitous generation of an EBV+ B-lymphoma xenograft from an NPC patient without clinically manifest or documented B cell lymphoma prompted us to explore whether the tumor developed from a pre-existing but innocuous and quiescent population of EBV-driven B-lymphoma cells, or via a *de novo* process during xenotransplantation. To investigate the former scenario, we re-examined 4-μm sections of the original NPC biopsy and dissociated cells which were cytospun onto slides. EBER-ISH signals were exclusively detected on pan-cytokeratin cells and absent in CD45+ lymphocytes, indicating that intratumoral B cells in the NPC biopsy were not likely to be infected with the virus (**Figure 2A**).

**Figure 2.**
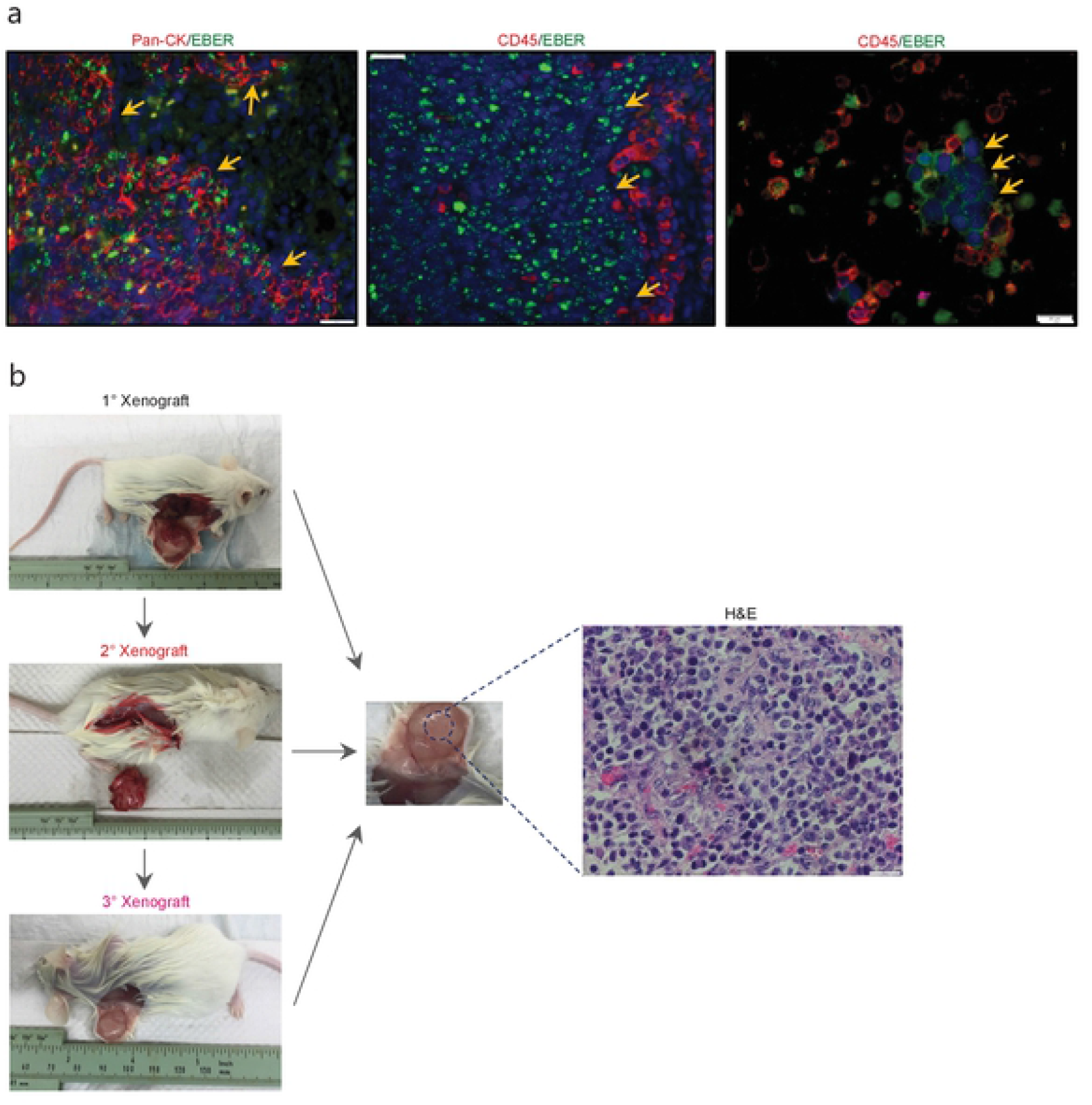
Histological characterization of the newly-established cell line. (a) To investigate whether the serendipitous generation of a B-lymphoma xenograft derived from a pre-existing but innocuous and quiescent population of EBV-driven B-lymphoma cells, we re-examined 4-μm sections of the original NPC biopsy and dissociated cells which were cytospun onto slides. EBERISH signals were exclusively detected on pan-cytokeratin cells and absent in CD45+ lymphocytes, suggesting that intratumoral B cells in the NPC biopsy were not likely to be infected with the virus. (b) Yu103 cells were tumorigenic when injected into NSG mice subcutaneously. The xenografts were serially passageable in mice and retained identical histology and morphology as the parental xenograft.

### Derivation and characterization of a new EBV-positive human B-cell lymphoma cell line

A new cell line, designated Yu103, was established from the xenograft model. Giemsa staining revealed a large B-cell lymphoma morphology (**Figure 3A**). Yu103 B-lymphoma cells can be passaged by gentle mechanical dissociation and re-plating onto RPMI1640 with 10% FBS without supplementation of growth factors, with a doubling time of 4-5 days. Cells also proved amenable to cryopreservation and remained viable after thawing.

**Figure 3.**
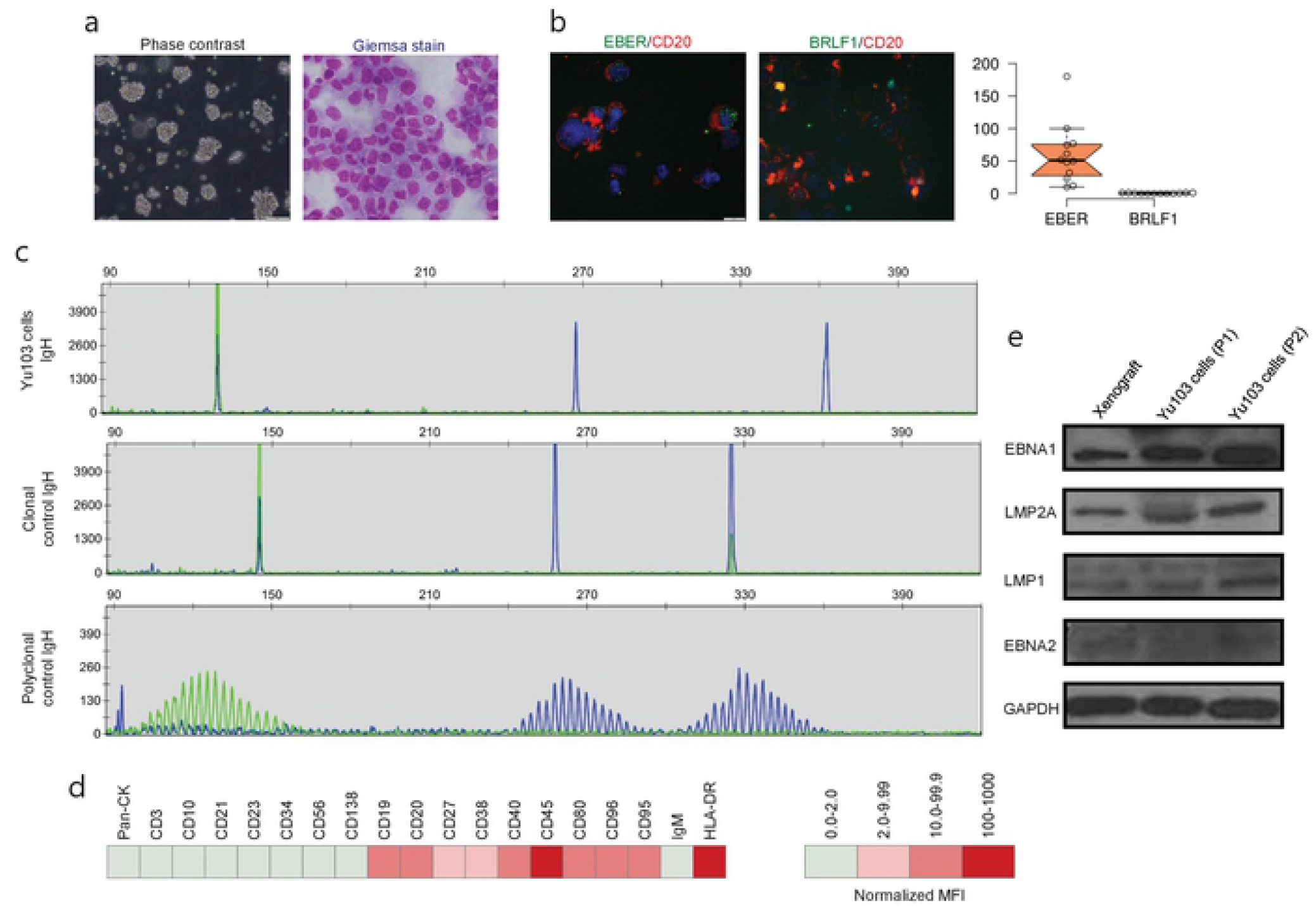
Molecular characterization of the Yu103 cell line. (a) Yu103 cells growing in free-floating clusters, and Wright-Giemsa staining shows large B-lymphocytes with visible nucleoli in the cytospun cell preparation. (b) Co-staining by immunocytochemistry of CD20 and *in situ* hybridization of EBER or BRLF1 in Yu103 cells. Quantitative estimates of EBV RNA puncta were plotted. (c) Yu103 cells were confirmed to be of monoclonal origin by PCR analysis of IgH gene rearrangements, which yielded single bands for the IgH genes centered around 130, 267 and 362bp respectively. The middle and lower panels show IgH analysis of monoclonal and polyclonal controls. (d) Fluorescence-activated cell sorting analysis and quantitative analysis of cell surface markers of Yu103 cells. (e) Western blot analysis of EBV latent gene products in parental xenograft tissues and passaged Yu103 cells (loading control: GAPDH)

Yu103 cells featured nuclear localization of biphasic EBV and characteristic CD20 expression on cell members as confirmed by fluorescent RNA *in situ* hybridization[19] and immunohistochemical staining respectively (**Figure 3B**). By visualizing and quantifying the punctate fluorescent signals, which correspond to EBV episomes, latent infection was found to be predominant (**Figure 3B**). Importantly, Yu103 cells were tumorigenic when injected into NSG mice subcutaneously. The xenografts were serially passageable in mice and retained identical histology and morphology as the parental xenograft (**Figure 2B**).

PCR analysis of the Yu103 cells for IgH gene rearrangements yielded a single band for the IgH gene, indicating a monoclonal origin (**Figure 3C**). Immunophenotyping of Yu103 cells by flow cytometry further demonstrated that they were positive for CD19, CD20, CD27, CD38, CD40, CD45, CD80, CD86, CD95 and negative for pan-CK,CD3, CD10, CD21, CD23, CD34, CD56 and CD138 markers (**Figure 3D**). Of note, the expression of the CD27 antigen indicates that Yu103 cells are of memory B cell origin[20]. Western blot assays performed on all samples from the xenograft to cell line revealed that Yu103 cells expressed the full repertoire of latent proteins that constitute the latency III program[21] (namely, EBNA1, EBNA2, LMP1 and LMP2A) (**Figure 3E**).

### Chemical and biological inducers promote entrance into the EBV lytic cycle

EBV-harboring cell lines whose transit from latency to the productive replicative cycle are chemically or biologically tractable are advantageous model systems for a variety of research applications[22–25]. Thus, we asked if EBV activation can be enhanced in Yu103 cells using common chemical (a combination of phorbol 12-myristate 13-acetate [PMA] and sodium butyrate) and biological (immunoglobulin [IgG] cross-linking of B-cell receptors) strategies. The EBV+ marmoset cell line B95-8 was used as a positive control and complexes formed between monoclonal BZ1 and the viral immediate-early protein ZEBRA (encoded by BZLF1), a sensitive and early marker for entry into the lytic phase, were quantified. Prior to stimulation, ~6% of Yu103 cells and 16% of B95-8 cells exhibited spontaneous replication (**Figure 4A and 4B**). At 48 h after induction, lytic replication in Yu103 cells non-significantly increased to ~7% with surface IgG cross-linking and to ~13% (*P* < 0.0001) with PMA plus sodium butyrate treatment.

**Figure 4.**
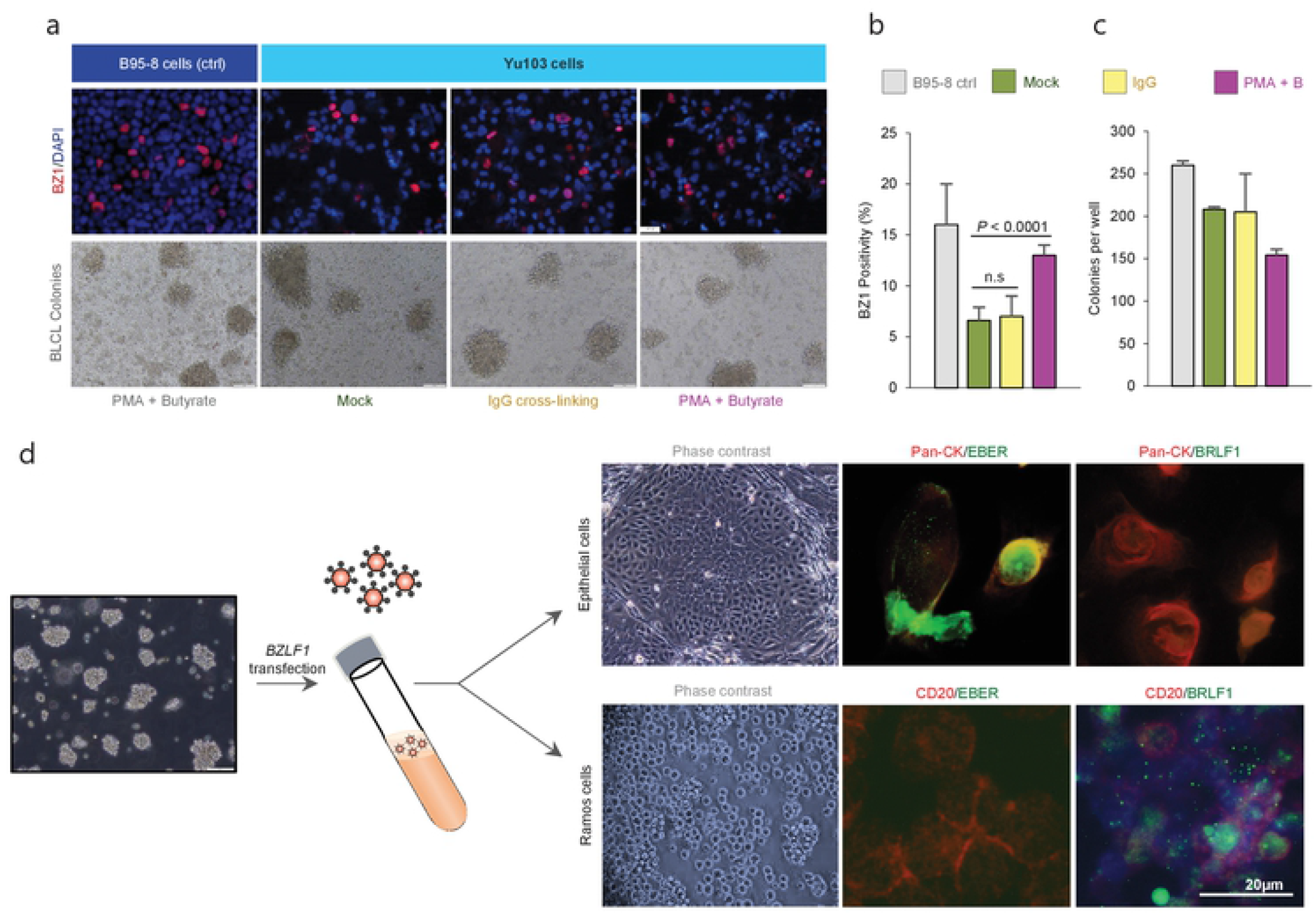
Tractability and cellular tropism of Yu103 EBV cells. (a-c) The amenability of Yu103 EBV to common chemical (a combination of phorbol 12-myristate 13-acetate [PMA] and sodium butyrate) and biological (immunoglobulin [IgG] cross-linking of B-cell receptors) induction methods were tested. We quantified complexes formed between monoclonal BZ1 and the viral immediate-early protein ZEBRA (encoded by BZLF1), a sensitive and early marker for entry into the lytic phase, and the EBV+ marmoset cell line B95-8 was used as a positive control. (d) Despite originating from nasopharyngeal carcinoma, Yu103 EBV were found to have comparable infective and transforming powers as B95-8 EBV, while retaining epitheliotropic abilities. Shown are representative panels of dual EBER-ISH (green) and immunocytochemistry of pan-CK (epithelial cells in red) or CD45 (lymphocytes in red) in consecutive sections at thickness of 4μm.

### EBV exhibits B-lymphotropism and immortalizes naïve B cells

In a *bona fide*, fully lytic replication, EBV generates prodigious amounts of virions which are capable of infecting and potentially immortalizing naïve B cells. To test this property of Yu103 EBV, denuded cell culture media containing a reservoir of Yu103 EBV virions was added to a milieu of peripheral blood mononuclear cells (PBMC) from a healthy EBV-seronegative donor. B95-8 EBV was used as a positive control under the same experimental set-up. Immortal PBMC colonies formed in the cell culture media contaminated with Yu103 or B95-8 EBV virions, while cells admixed with control cell culture media perished. By quantitating the rate of immortalized B-lymphoblastoid cell lines (BLCL) colony formation, Yu103 EBV were found to have comparable infective and transforming powers as B95-8 EBV (**Figure 4C**).

### Cell-free Yu103 EBV retains epitheliotropism for nasopharyngeal epithelium

While most EBV strains associated with malignancies are avidly B-lymphotropic, they are also extremely inefficient at infecting epithelial cells except for the few strains linked to nasopharyngeal carcinoma [5,7–9,26]. Cell-free infection of epithelial cells is even more challenging and has only been exemplified by strains with an exceptional degree of epitheliotropism [5,8,9,27], such as the recently-isolated M81 strain from a Chinese male with nasopharyngeal carcinoma[8]. Since Yu103 EBV progenitors originated from nasopharyngeal carcinoma cells, a valid enquiry is whether Yu103 EBV progeny – which currently propagate in B-lymphoma cells – still retain the proclivity for infecting and replicating in nasopharyngeal epithelial cells.

To investigate the epitheliotropism of Yu103 EBV, we induced virion production via BZLF1 plasmid transfection and added genomic DNA-equivalent cell-free supernatant to primary nasopharyngeal epithelial cells and EBV-negative Ramos cells. Infection occurred in both cell types with distinct patterns: epithelial cells primarily exhibited EBER+ latent infection, while lytic infection prevailed in Ramos cells (>90%) (**Figure 4D**). As such, these experiments collectively demonstrate that Yu103 EBV possesses an uncanny and distinctive trait of being simultaneously epitheliotropic and B-lymphotropic.

### Whole-genome sequencing and miRNA microarray profiling of Yu103 EBV

One possible scenario which might account for some of the experimental results obtained thus far, but which has not been formally tested in the preceding investigations, is the notion that the B cell lymphoma PDX and patient’s original nasopharyngeal carcinoma tumor were driven by distinct strains of EBV which simultaneously infected the source patient. To our knowledge, coinfection of the same individual by two *different* clonal EBV strains, each associated with either nasopharyngeal carcinoma or B-lymphoma, has only been documented in one case report of a patient who developed both malignancies [28], indicating that such a scenario is plausible albeit vanishingly unlikely.

We therefore proceeded to perform whole-genome sequencing on DNA samples extracted from the original patient tumor biopsy, xenograft, and Yu103 cells. Based on the coverage of the target regions, DNA sequences in all three samples bared most resemblance to the HKNPC1 cell line, which was subsequently used as the reference genome for *de novo* assembly. Sequencing achieved a target coverage of 92.6% to 96.1% and an average depth of 3,480 × in all three samples, in spite of bioinformatic challenges posed by the low abundance of viral DNA relative to human DNA and internal repeat sequences.

Our 3 new sequences were then compared with each other as well as 19 other published EBV sequences [29,30]. We inferred evolutionary histories by computing maximum composite likelihood estimates of the evolutionary distances[31], which were subjected to the neighborjoining algorithm to reconstruct the topology of the most statistically-parsimonious phylogenetic tree [32] (**Figure 5A**). Yu103 EBV was found to cluster in a clade with other Type 1 strains (which are defined by polymorphisms in the EBV nuclear antigens, *EBNA-2*, −3, −4, −6), but not AG876, the only type 2 EBV strain derived from a Ghanaian patient with Burkitt’s lymphoma (**Figure 5B**). Not surprisingly, Yu103 was closely related to Asian nasopharyngeal carcinoma isolates such as C666-1, M81, HKNPC1 and GD2, but distant from wild type, B-lymphoblastoid, and B-lymphoma strains (e.g. WT4, K4123-MiEBV, K4413-MiEBV, Akata, and B95-8).

**Figure 5.**
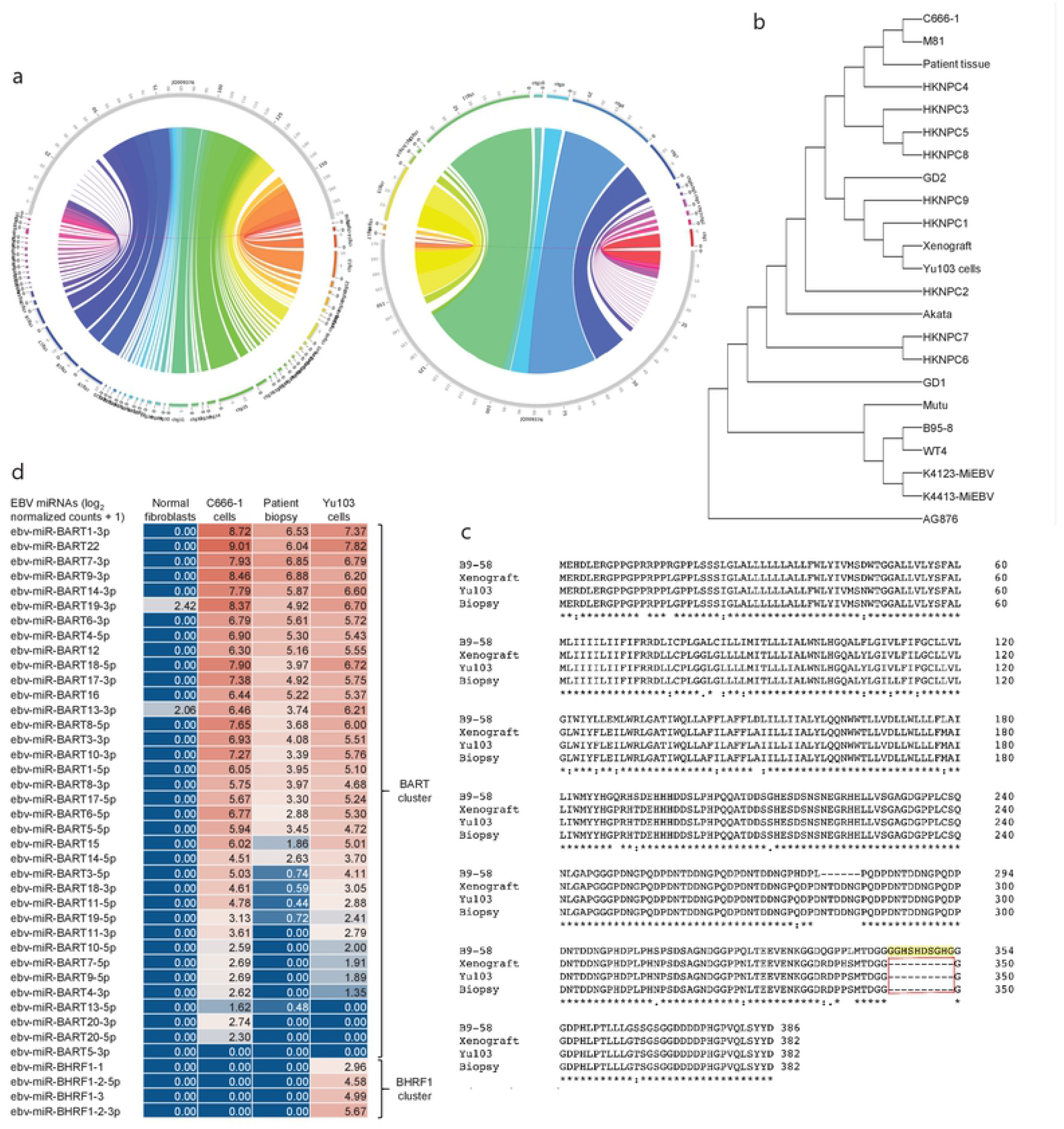
Molecular characterization and phylogenetic analysis of Yu103 EBV. (a-b) Neighbor-joining phylogenetic tree depicting the genomic relatedness of EBV strains based on whole-genome sequencing with maximum composite likelihood. We compared n=23 EBV isolates including Yu103 cells (original patient tissue, xenograft, and purified cell line), NPC strains (M81, C666, HKNPC, GD1, GD2), gastric cancer strains and lymphoma strains (Raji, AG876 and Akata) All positions containing gaps and missing data were eliminated, with a total of 123852 positions in the final dataset. (c) Amino acid sequence variation between the variant LMP1 genes derived from B95-8, Yu103 cell, xenograft and original NPC biopsy. Asterisks (*) indicate positions with a single, fully conserved residue; colons (:) indicate strong conservation with scores of >0.5 in the Gonnet PAM 250 matrix; periods (.) indicate weak conservation with scores of ≤0.5 in the Gonnet PAM 250 matrix; and dashes (-) represent deleted amino acids. Yu103 EBV from the Singapore Chinese male in this study was found to harbor a 30-base pair deletion in *LMP1* (observed in the original patient biopsy, xenograft, as well as isolated Yu103 cells) which is known to occur at a high prevalence among EBV strains isolated from Hong Kong Chinese patients with undifferentiated nasopharyngeal carcinoma. (d) Microarray-based miRNA scrutiny demonstrated that Yu103 EBV shared a similar BamHI A Rightward Transcripts (BARTs) miRNA expression profile as that of the prototypical EBV-positive NPC cell line, C666-1. However, a major distinction was that a cluster of miRNAs which regulate *BHRF1*, the viral homologue of the Bcl-2 protooncogene, was exclusively expressed by Yu103 but not C666-1 EBV, which may account for Yu103’s peculiar ability to efficiently infect and subsequently induce lymphomagenesis in B cells. miRNA expression were analyzed on a log_2_ [normalized counts + 1] scale.

Next, we interrogated amino acid sequences of specific candidate genes known or postulated to mediate the tropism and virulence of EBV (**Figure 5C**). We honed in on the latent membrane protein 1 *(LMP1)* oncogene because it has a well-established role in EBV-driven carcinogenesis and other aspects of their cancer biology such as immunosurveillance [15,33,34], and has been shown to harbor polymorphisms which may confer a degree of epitheliotropic predilection in EBV strains associated with nasopharyngeal carcinoma[5,8,9]. Yu103 EBV from the Singapore Chinese male in this study was found to harbor a 30-base pair deletion in *LMP1* which is known to occur at a high prevalence among EBV strains isolated from Hong Kong Chinese patients with undifferentiated nasopharyngeal carcinoma[35–38] (**Figure 5B**). Microarray-based miRNA scrutiny demonstrated that Yu103 EBV shared a similar BamHI A Rightward Transcripts (BARTs) miRNA expression profile as that of the prototypical EBV-positive NPC cell line, C666-1 (**Figure 5D**). However, a major distinction was that a cluster of miRNAs which regulate *BHRF1*, the viral homologue of the Bcl-2 proto-oncogene, was exclusively expressed by Yu103 but not C666-1 EBV, which may account for Yu103’s peculiar ability to efficiently infect and subsequently induce lymphomagenesis in B cells [19,39–41] (**Figure 5D**).

## DISCUSSION

The Epstein-Barr virus (EBV) is responsible for 1-2% of the global cancer burden, or approximately ~200,000 new cases of malignancies each year [16]. Development of effective therapies and prophylactic strategies against EBV-driven neoplasia have been severely constrained by our inability to recapitulate the diversity of pathogenic EBV strains in preclinical research, which underscores the importance of expanding our armamentarium of experimental tools. In this study, we report the discovery and characterization of a newly-identified Asian Pacific EBV strain (Yu103), which has a remarkable provenance of being isolated from a nasopharyngeal carcinoma (NPC) biopsy but subsequently propagating in human B-lymphoma cells where it was found to exhibit unusually high levels of spontaneous lytic reactivation. As such, we envision that the unique properties of Yu103 EBV, as well as the new B-lymphoma cell line and xenografts it is stably established in, will provide a valuable and unprecedented platform for further unravelling the pathogenetic mechanisms of EBV oncogenesis and facilitating the development of novel clinical modalities.

Unlike most hitherto characterized oncogenic EBV strains which have been described to exclusively drive either lymphomagenesis or carcinogenesis, Yu103 evinces an uncanny association with both nasopharyngeal carcinoma (NPC) and B-cell lymphoma. Indeed, one of the most enigmatic aspect of EBV has been the epidemiological phenomenon that although it homogenously infects human adult populations from all geographic regions, the incidence of neoplastic diseases which it spawns is extremely skewed [2,3]. Another phenomenon widely appreciated by clinicians is the fact that EBV-associated NPC and lymphomas of the head and neck region rarely ever afflict the same individual, with no more than five case reports in the literature documenting the synchronous or sequential diagnosis of both types of neoplasms [28,42–44]. In the only case report in which viral genotyping was performed, it was revealed that the patient’s (Hodgkin) lymphoma and NPC were driven by distinct EBV strains[28]. Such observations have consequently fostered the hypothesis that divergent strains of EBV exist worldwide with very contrasting predilections for infecting, replicating in, and transforming epithelial or lymphocytic cells[8]. The discovery and characterization of Yu103 offers a counterexample to such a proposition.

There is merit in the notion that the serendipitous establishment of an EBV-positive human B-lymphoma xenograft represents the outgrowth of an elusive and quiescent population of EBV-driven B-lymphoma clones residing within the NPC biopsy, which gained a competitive advantage in the immunodeficient mice. Moreover, if B-lymphoma cells were already pre-existent within the NPC biopsy, it is certainly plausible that they could also have been transformed by a distinct EBV strain compared to those driving NPC cells, as has been documented in a previous case report [28]. These scenarios are, however, unlikely because of several countervailing observations. Firstly, the patient’s presentation and physical examination was not consistent with clinically-manifest B-lymphoma, and such a diagnosis was further excluded during the multiple sets of radiological and histopathological investigations ordered as part of routine clinical care. Secondly, histopathological re-examination of the original NPC biopsy by a board-certified pathologist also failed to detect the presence of any intratumoral B-lymphoma cells. Thirdly, although two natural reservoirs of EBV exists in healthy asymptomatic carriers (i.e., cell-free EBV in saliva or cellular EBV trafficking with latently infected memory B cells as they circulate through the body and back to Waldeyer’s ring [45]), our evolutionary tree analyses indicated that the three isolates in this study were distant from the wild-type saliva EBV (WT4) and expressed CD27, pointing to their memory B cell origin.

Fourthly, we also re-examined sections of the NPC biopsy using RNAscope *in situ* hybridization assays with single-molecule sensitivity for detecting EBV-specific gene products [17–19], and observed that EBER-ISH signals were exclusively detected on pan-cytokeratin cells but absent on CD45+ lymphocytes. Finally, by parsing the whole-genomes of EBV isolates from the original NPC biopsy, xenograft, and cell line, we were able to ascertain that these three EBV isolates exhibit homologous sequences at highly discriminative genomic loci such as the genes encoding the nuclear antigens *(EBNA-2*, −3, −4, −6) and *LMP1*. These observations therefore suggest that the emergence of a B-lymphoma colony is more likely to represent a *de novo* process in which the same EBV strain found in NPC cells infected and subsequently induced malignant transformation of memory B-lymphocytes during xenotransplantation, rather than the preferential outgrowth of an obscure population of B-lymphoma cells driven by a discrete B-lymphotropic EBV strain.

Of greater translational relevance are the biological properties that endow Yu103 EBV its distinctive capacity to drive both NPC and B-lymphoma. The ability of EBV to target and infect epithelial cells is the foremost requirement – and invariably the rate-limiting step – in the multistep process of EBV-driven carcinogenesis. Although most EBV strains characterized in the context of malignancies are known to be avidly B-lymphotropic, infection of epithelial cells *in vitro* proves far more challenging, but can be enhanced by co-culture with Akata cells expressing recombinant EBV presumably via cell-to-cell contact [5,8,9,27]. The notion that epithelial cells are rarely targeted by EBV is also exemplified in *in vivo* studies which have shown that latent EBV infection is absent in nasopharyngeal tissue biopsies obtained from chronically-infected individuals, indicating that the virus is not a regular passenger in the nasopharyngeal epithelium[2]. Not surprisingly, Yu103 EBV evinced the ability to infect primary nasopharyngeal epithelial cells even when cells were admixed with cell-free viral supernatant – a defining feature of epitheliotropic viral strains linked to carcinoma formation [5,8,9,27]. After infecting epithelial cells, EBV strains associated with carcinomas must also be able to commandeer the molecular machinery in host cells in order to engender malignant transformation. To investigate Yu103’s carcinogenic potential, we honed in on the well-established EBV oncogene, latent membrane protein 1 *(LMP1)*, which functions as a constitutively active tumor necrosis factor receptor (TNFR) engaging a myriad of downstream anti-apoptotic and proliferative signaling cascades[15,33,34]. Interestingly, Yu103 EBV was found to harbor a 30-bp deletion polymorphism in the third exon of *LMP1*, which encodes a 10-amino acid sequence within the C-terminus activation region (CTAR)-2 domain of the protein. This particular genetic alteration promotes the retention of EBV genomes in latently-infected cells and enhances the carcinogenic potential of the virus[15,33,34], and is known to occur at a disproportionately high prevalence in geographical regions where NPC is endemic [35–38]. Another salient finding is the overexpression of BamHI A Rightward Transcripts (BARTs) miRNAs in Yu103 cells, which intriguingly, was found to mirror the expression profile of the prototypical EBV-positive NPC cell line, C666-1. This observation probably reflects Yu103 EBV’s provenance from NPC but may also account for its epithelial cell-transforming powers, since the targets of BART miRNAs include LMP1 and other putative gene products that may be critical to preventing the eviction of the EBV episomal genome from host epithelial cells[46,47].

The inception of a new B-lymphoma cell line and PDX model likewise warrants discussion. Since Yu103 EBV’s ability to infect and propagate in B cells is a known contrivance of all EBV strains (all of which establish a life-long carriage in the host by targeting and propagating in memory or resting B cells), a more pertinent area of enquiry are the mechanisms responsible for malignant transformation after Yu103 gained entry into memory B cells. One plausible mechanism is suggested by the miRNA screen and relates to the observation that a cluster of miRNAs which regulate *BHRF1* (the viral homologue of the Bcl-2 proto-oncogene) was neither expressed in the original NPC biopsy nor in the C666-1 NPC cell line but were highly expressed in Yu103 B-lymphoma cells. This is consistent with previous studies which suggest that the overexpression of BHRF1 miRNAs (eg. miR-BHRF1-2 and miR-BHRF1-3) serves to license or facilitate the latency III program and productive lytic cycle that characterizes many primary EBV-driven lymphomas [19,39–41]. Indeed, in consonance with their overexpression of EBV-encoded BHRF1 miRNAs, Yu103 cells were found to undergo spontaneous lytic reactivation at unusually high levels and express the full repertoire of latent proteins that constitute the latency III program[21]. As such, these shifts in miRNA transcriptomes could represent a novel molecular switch exploited by Yu103 to alter between the predominantly-latent replicative phase in NPC and the more productive lytic cycle in B-lymphoma cells.

The discovery of Yu103 EBV and its establishment in a new B-lymphoma cell line and xenograft will provide valuable resources for research on EBV-associated hematological and epithelial malignancies. Although a handful of *recombinant* EBV strains (e.g. M81 and ABA) have since been established from NPC cells and show similar properties to Yu103, such as the ability to undergo spontaneous lytic replication in lymphoblastoid cells when admixed with adult primary B cells, an important distinction is that these *recombinant* strains were expressly engineered to acquire pathogenic capabilities, whereas Yu103 was directly established without being subjected to any genetic manipulations [8]. The newly-established Yu103 cell line in our study has a high mitotic index and rapid doubling time; can be passaged by gentle mechanical dissociation and replating without supplementation of growth factors; readily forms xenograft tumors with large B-cell lymphoma morphology when implanted in immunodeficient mice; and is amenable to chemical induction strategies such as treatment with PMA and sodium butyrate [22–25]. For future research, it will be instructive to perform integrated multidimensional analyses of Yu103 EBV and the host cells it resides in; probing the antigenicity and immunogenicity of Yu103 EBV-driven tumors, and appraising the implications for EBV-directed cancer immunotherapies [2,14–16]; and investigating whether Yu103 is capable of transforming gastric epithelium, since the LMP1 deletion variant which it harbors has been detected in ~80% of EBV-associated gastric cancer cases in Asia [48,49].

## METHODS

### Ethics Statement

All animal protocols conformed with the guidelines set forth by the Institutional Animal Care and Use Committee of the National University of Singapore, who approved the experimental protocol (protocol R14-0144). All assays utilizing human samples were reviewed and approved by the National Healthcare Group Domain-Specific Review Board, and informed and written consent was obtained from the patient in our study.

### NPC specimen

A biopsy was obtained from a 38-year-old Singaporean Chinese male with stage 4 (T3N1M0) nasopharyngeal carcinoma. A portion of the biopsy was submitted to the pathology laboratory for diagnosis and the remaining portion was immediately immersed and extensively washed in a Hank’s balanced salt solution containing penicillin, streptomycin, and amphotericin B. A small piece (~1 mm^3^) of the specimen was digested with 10 mg/ml of Dispase II (Sigma, St. Louis, MO) at 4°C overnight, followed by dissociation by repetitive pipetting; while the remainder unused specimen was fixed. Enzymatic digestion was stopped by adding media containing serum. The dissociated cells were explanted onto F-culture media as previously described[50].

### Xenograft model

Four-week-old NOD-SCID/Il2rg2/2 (NSG) mice were subcutaneously injected with P1 cell cultures derived from the NPC biopsy. Mice were monitored for changes in bodyweight and tumor burden weekly. Mice were sacrificed when tumor size reached a diameter of 1.5 cm.

### Histology and staining

Formalin-fixed, paraffin-embedded tissue were sectioned to 4 um thickness for IHC and immunofluorescent staining as previously described [51]. Antibodies were purchased from Abcam (Cambridge, UK) and used at 1:100~200 dilutions. *In situ* hybridization was performed according to the protocol provided by the manufacturer (Advanced Cell Diagnostics; Newark, USA). For Giemsa staining, dissociated cells were cytospun onto slides was fixed in absolute methanol for 30 seconds, and then submerged in 10% Giemsa solution for 30 minutes. Fluorescent *in situ* hybridization of EBV viral mRNA with immunohistochemical (IHC) staining was performed as described in our previous study[17]. All images were captured using an Olympus fluorescent microscope equipped with appropriate filters, and image analysis was conducted using the in-built Cellsens software.

### Cell culture

Yu103 cells were dissociated from resultant mouse xenograft, and initially cultured in RPMI-1640 media supplemented with 20% fetal bovine serum (FBS) at 37°C with 5% CO2. When constant cell growth was achieved, the culture medium was replaced with RPMI-1640 supplemented with 10% FBS. Passaging was performed through gentle pipetting. B95-8, Akata, and Ramos cells were cultured in RPMI-1640 media supplemented with 10% FBS. Primary nasopharyngeal epithelial cells were cultured in F media as previously described [50].

### BLCL immortalization

To assess the rate of immortalized B-lymphoblastoid cell line (BLCL) colony formation, PBMCs were resuspended to a density of 2 × 10^6^ cells/ml and admixed with an equal volume of denuded cell culture medium harvested from induced Yu103 cells. The mixture was then incubated at 37°C with 5% CO_2_ incubator for 2 h, washed and resuspended to a final density of 1 × 10^6^ cells/ml using culture medium supplemented with 1 μg/ml cyclosporin A (Sigma-Aldrich). Cells were incubated for three weeks in culture media supplemented with cyclosporin A, and the formation of macroscopic cell clusters was quantified thereafter.

### Immunophenotyping

5 × 10^5^ Yu103 cells were resuspended in flow cytometry buffer (1 × phosphate-buffered saline supplemented with 3% fetal bovine serum and 0.01% sodium azide) and incubated with fluorophore-conjugated antibodies. For typing of cell surface lineage markers, cells were stained for pan-CK (AE1/AE3) (eBioscience), CD3 (HIT3a), CD10 (HI10a), CD21 (LT21), CD23 (EBVCS-5), CD34 (581), CD56 (HCD56), CD138 (1D4), CD19 (HIB19), CD20 (2H7), CD27 (O323), CD38 (HB-7), CD40 (G28.5), CD45 (HI30), CD80 (2D10), CD86 (IT2.2), CD95 (DX2), IgM (MHM-88) and HLA-DR (L243). All antibodies were purchased from BioLegend unless stated otherwise. Samples were acquired on BD LSRFortessaTM (Becton Dickinson) and analyzed using FlowJo software v7.6.5 (TreeStar). For the purpose of illustration and categorization in Figure 3, normalized median fluorescence intensity (nMFI) values of 0 to less than 2 indicates negative staining (-), above 2 to 9.99 (+), 10 to 99.9 (++) and 100 to 1000 (+++).

### Western blot

Cells were lysed in RIPA buffer containing complete mini protease inhibitor (Roche) for 15 minutes on ice with occasional vortexing. Samples were centrifuged, and cell lysates were mixed with SDS loading buffer and boiled at 95°C for 10 minutes. Sample were then loaded and separated on 10% SDS-PAGE gels, before transferring onto polyvinylidene difluoride (PVDF) membranes. Membranes were blocked with 5% milk diluted in PBS containing 0.05% Tween-20 (PBST) for 1 hour, before incubation with primary antibodies at 1:500-1000 dilution overnight at 4°C. Thereafter, membranes were washed and probed with horseradish peroxidase (HRP)-conjugated secondary antibodies for 1 hour before detection with Western Lightning Chemiluminescence (Perkin Elmer). Antibodies used included: anti-EBNA1 (sc-57719, Santa Cruz), anti-EBNA2 (ab90543, Abcam), anti-LMP1 (CS1-4, Dako), anti-LMP2A (MCA2467, Biorad), anti-GAPDH (mab374, Merck), anti-mouse IgG-HRP (ThermoFisher Scientific) and anti-rat IgG HRP (Santa Cruz).

### EBV induction and cell-free viral infection

Yu103 cells were treated with a combination of 20 ng/ml sodium butyrate (SB) plus 3 mM phorbol 12-myristate 13-acetate (PMA), and incubated for 48 h. The supernatant was collected by centrifugation and filtered through 0.45μm syringe filter to ensure that the supernatant was devoid of cells. Titration of cell-free virions was done by real-time PCR using an EBNA1 Taqman probe. After digestion with DNase I, EBV genomes were quantified based on the standard curve generated from 10-fold serial dilutions of genomic DNA of Namalwa cells, which have two integrated copies of the EBV. Two milliliters of cell-free supernatant was admixed with 2 × 10^5^ primary nasopharyngeal epithelial cells or Ramos cells in six-well plates in 2 ml of medium (~7 × 10^7^ DNA copies/ml). Cells were fixed after 24 h and cell-free infection was assessed by EBERISH and BRLF1-ISH as previously described [17].

### EBV whole-genome sequencing

Total genomic DNAs (containing human and viral genomes) were isolated using AllPrep DNA/RNA Mini Kit (Qiagen) from the NPC biopsy, cell line, and xenograft, and samples were submitted to MyGenostics (Beijing, China) for next-generation sequencing (NGS) analysis. DNA library preparation, probe designs, target capture using *in situ* hybridization, index tagging, sequencing, de novo assembly, and bioinformatics analysis were performed using published protocols which have been optimized and validated by the manufacturer for NGS of the EBV genome [30,52–54]. We inferred evolutionary histories by computing maximum composite likelihood estimates of the evolutionary distances[31], which were subjected to the neighborjoining algorithm to reconstruct the topology of the most statistically-parsimonious phylogenetic tree [32] using a total of 23 nucleotide sequences [29,30]. All positions containing gaps and missing data were excluded, and a total of 123,852 positions were used in the final dataset. Phylogenetic analyses were conducted in MEGA7 [55].

### MicroRNA array

miRNA was isolated from the Yu103 cell line, NPC biopsy, cancer-associated fibroblasts (CAFs) and C666-1 cells using RNeasy Mini kit per the manufacturer’s instructions (Qiagen, Hilden, Germany). RNA integrity number (RIN) was ascertained using Agilent 2100 Bioanalyzer (Agilent Technologies Inc., USA). Hybridization and scanning were performed according to the Agilent miRNA protocol using the miRNA 3.0 array (AMADID 085021), which covers 2,589 probes for human (2549) and EBV (40) miRNA.

### Lead Contact and Materials Availability

There are restrictions to the availability of Yu103 xenograft and cell lines due to the lack of an external centralized repository for its distribution and our need to maintain the stock. We are glad to share these materials with reasonable compensation by the requestor for its processing and shipping.

